# Reward-driven adaptation of movements requires strong recurrent basal ganglia-cortical loops

**DOI:** 10.1101/2024.10.15.618421

**Authors:** Arthur Leblois, Thomas Boraud, David Hansel

**Author notes:** Corresponding author; Arthur Leblois, **Email :**.

## Abstract

The basal ganglia (BG) are a collection of subcortical nuclei involved in motor control, sensorimotor integration, and procedural learning. They play a key role in the acquisition and adaptation of movements, a process driven by dopamine-dependent plasticity at cortico-striatal projections, which serve as BG input. However, BG output is not necessary for executing many well-learned movements. This raises a fundamental question: how can plasticity at BG input contribute to the acquisition and adaptation of movements which execution does not require BG output? Existing models of BG function often neglect the feedback dynamics within cortico-BG-thalamo-cortical circuitry and do not capture the interaction between the cortex and BG in movement generation and adaptation. In this work, we address the above question in a theoretical model of the BG-thalamo-cortical multiregional network, incorporating anatomical, physiological, and behavioral evidence. We examine how its dynamics influence the execution and reward-based adaptation of reaching movements. We demonstrate how the BG-thalamo-cortical network can shape cortical motor output through the combination of three mechanisms: (i) the diverse dynamics emerging from its closed-loop architecture, (ii) attractor dynamics driven by recurrent cortical connections, and (iii) reinforcement learning via dopamine-dependent cortico-striatal plasticity. Our study highlights the role of the cortico-BG-thalamo-cortical feedback in efficient visuomotor adaptation. It also suggests a mechanism for early-stage acquisition of reaching movements through motor babbling. More generally, our model explains how the BG-cortical network refines motor output through its intricate closed-loop dynamics and dopamine-dependent plasticity at cortico-striatal synapses.

**Significance statement:** The basal ganglia (BG) are essential for learning and adapting movements, yet their output is often unnecessary for executing well-learned actions. This paradox challenges existing models of BG function. Here, we present a biologically grounded model of the cortico-BG-thalamo-cortical loop, showing how this closed-loop network supports efficient motor adaptation and early movement acquisition. Our findings reveal that dopamine-driven plasticity, recurrent cortical dynamics, and feedback from the BG together shape motor output, even when BG output is not directly required for execution. This work provides a unified framework explaining how the BG-cortical circuit contributes to both learning and refinement of goal-directed actions.

## Introduction

The acquisition of motor skills relies on large brain networks involving, among others, motor cortical areas and the basal ganglia (BG), a set of deep brain nuclei. The BG play a key role in sensorimotor integration and procedural learning (1–3). They are also essential for reinforcement-driven adjustments during the acquisition of motor skills (1, 4). Notably, optogenetic modulation of dopaminergic input to the BG can drive sensorimotor adaptation (5). In mechanistic circuit models of reinforcement learning (RL), adaptation relies on dopamine-driven long-term plasticity at cortico-striatal synapses (3, 6, 7). According to these models, reward-based learning alters these synapses, modifying cortical input to the BG and refining their output to optimize reward-driven behavior (3).

However, experimental evidence indicates that circuits outside the BG, such as the motor cortex, are the primary drive of motor output (8, 9). In humans, pallidotomy—a procedure that lesions the BG output nucleus—effectively reduces motor symptoms in Parkinson’s disease and in dystonia but with minimal negative impact on movement (10, 11). Pharmacological manipulations of the GABAergic system within the GPi in NHPs alter movement vigor and kinematic(12). Yet, it does not prevent the execution of well-practiced skills, suggesting that the BG do not store familiar or overlearned movements(13, 14).

During motor learning, neuronal activity initially changes in BG input structure, *i*.*e*. the striatum, and later in the motor cortex (15, 16). As learning progresses, behavioral adjustments become increasingly independent of the BG (17, 18), while gradual synaptic changes (19) and large-scale restructuring take place in the motor cortex (20). Current models of BG function do not explain how the BG can alter motor output during learning without being essential for the execution of well-practiced skills (14).

The cognitive, associative, motor, and premotor cortical areas provide significant input to the BG through their projections to the striatal and subthalamic regions (21). The BG-thalamo-cortical network is organized in parallel feedback loops at cellular spatial scale (22–24). This anatomical organization likely contributes to its normal and pathological dynamics (25–28). In this study, we develop a new theoretical model of the multiregional cortico-BG-thalamo-cortical network to explore its role in reward-based movement adaptation and how its dynamics influence the execution of reaching movements.

## Results

### The cortico-BG-thalamo-cortical circuit model

During reaching movement execution the response of the neurons in the motor cortex is tuned to specific movement parameters, *e*.*g*., direction or velocity (29). This fact led to the view that the motor cortex primarily encodes the movement parameters. A read out of this activity is performed in the spinal cord via a distributed population code(30). Similarly, neurons in the output nucleus of the BG exhibit tuning to movement features (31, 32). Building on this perspective, several theoretical studies have extended the “ring attractor framework” (33) to model the motor cortex, investigating how recurrent connectivity shapes movement-related neural activity in M1 (34).

Here, we develop a rate-based dynamical model of the basal ganglia–thalamo–cortical network within the ring attractor framework to investigate the role of the basal ganglia in reward-based adaptation of reaching movements. It encompasses seven neuronal populations (Fig. 1A): the striatum, the internal and external segments of the globus pallidus (GPi and GPe), the subthalamic nucleus (STN), the thalamus and two cortical populations. One cortical population represents M1 (M_ctx_) and the other one corresponds to the posterior parietal cortex (PCC), an associative cortical area. The latter encodes the location of sensory cues and is involved in the sensorimotor integration during reaching movement execution (35, 36).

**Figure 1:**
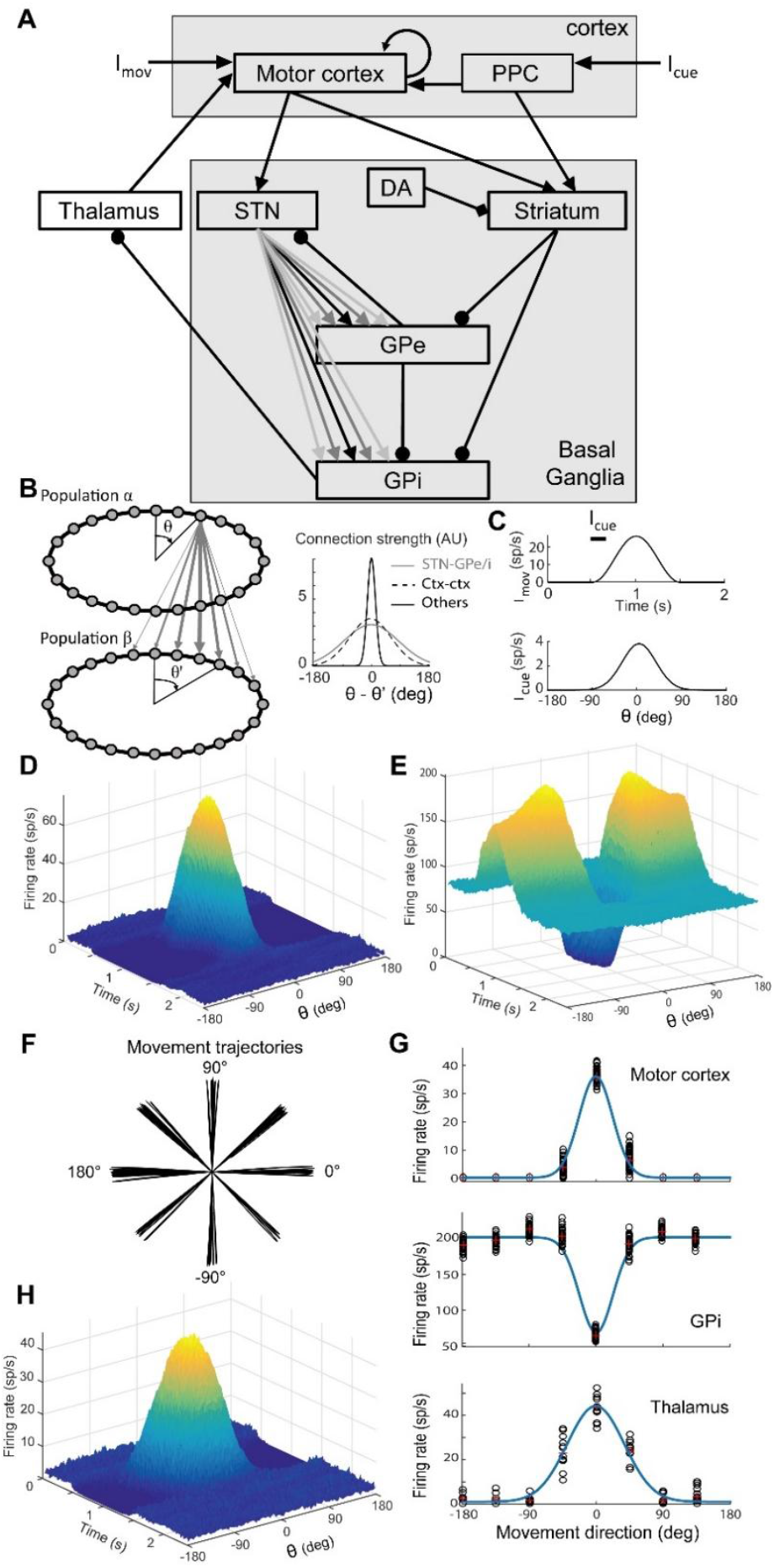
The BG-thalamo-cortical network model, tuning of synaptic weights, movement-related activity in the network and tuning curves in the cortex and GPi. A: Architecture of the BG model network. Each neural population is represented by a ring where neurons are evenly distributed and associated with a PD between - 180° and 180°. B: The connectivity between the different populations of the network is focal and organized in parallel beams (bottom right inset). In two connected populations, the strength of the connection between two neurons from these two populations is a gaussian function of the angular distance between the PDs of the two neurons (see Methods). The width of the gaussian connectivity function is tight (σ = 9°) in all connections of the network except for the divergent projection from STN to GPe and GPi (σ = 360°) and the recurrent connections in Mctx. C: Movement-related input in the model. Top: time course of the input **I**_**mov**_ added to the cortical population to induce movement. The thick black line indicates the duration of the weak and transient movement-related input to the PPC **I**_**cue**_ (pointing the direction of movement). Bottom: Profile of the weak and transient movement-related input to the striatum. D: Movement-related activity induced in the cortex in the presence of the full closed-loop BG-thalamo-cortical network. The cortical population displays a strong activity bump peaking at the angle pointed by the input from the PPC (**I**_**cue**_). E: the GPi population displays a sharp profile with decreased activity around the movement direction and increased activity away from the movement direction. F: Example trajectories generated in the model using a population vector computed as the sum of the PD of motor cortical neurons weighted by their firing rate at each time step. Note that the variability in direction and amplitude is due to the synaptic noise introduced in the input to neurons. G: Tuning curve of the movement-related activity for neurons in the cortex (Top) and GPi (Bottom). The angle associated with each neuron in the population corresponds to the preferred direction, as defined by the movement direction evoking maximal activity increase (in the cortex) or decrease (in the GPi). H: Movement-related activity induced in the cortex following the inhibition of the GPi (50% randomly selected neurons silenced in the GPi). Note that the cortical population still displays a strong activity bump peaking at the angle pointed by the input from the PPC (**I**_**cue**_), although with lower amplitude compared to the situation with intact BG-cortical loop in D. Parameters are as in Table 1 (see Methods).

The model’s architecture mirrors the anatomy (2, 21) and the connectivity (21, 22) of the motor loop within the cortico-basal ganglia-thalamo-cortical network. It includes three pathways from the cortex to BG output nucleus GPi: the direct pathway through the striatum, the indirect pathway through striatum, GPe, and STN, and the hyperdirect pathway through the STN (37). For simplicity, the various neuronal populations in the striatum participating in the direct and indirect pathways are merge into a single population (see Discussion).

**Table 1.** Values of the various parameters of the model. Spontaneous firing rates (m_0α_) are expressed in spikes/s, standard deviations (σ_αβ_) for connectivity patterns in degrees, delays (Δ_αβ_), time constants (τ_αβ_) and dt in ms, Other parameters are unitless. Note that J_str,Mctx_ and/or J_STN,Mctx_ may vary in various figures of the article. Finally, the external inputs to all populations (H_α_) are calculated as to lead to the activity levels displayed here as (m_0α_, in spikes/s) in the absence of movement-related input.

|  |  |  |  |  |  |
| --- | --- | --- | --- | --- | --- |
| $m_{0M_{ctx}}$ | 2 | $S_{M_{ctx}}$ | 0.02 | dt | 0.5 |
| $m_{0Str}$ | 0 | $S_{Str}$ | 0.02 | $\eta$ | $2 \times 10^{-5}$ |
| $m_{0STN}$ | 20 | $S_{STN}$ | 0.03 | $\beta$ | 4 |
| $m_{0GPe}$ | 60 | $S_{GPe}$ | 0.03 | $\sigma_c$ | 45 |
| $m_{0GPi}$ | 80 | $S_{GPi}$ | 0.05 | $s_c$ | 1 |
| $m_{0Tha}$ | 20 | $S_{Tha}$ | 0.02 | $\tau_c$ | 50 |
| $\Delta_{str,M_{ctx}}$ | 5 | $J_{str,M_{ctx}}$ | 2 | $\tau$ | 5 |
| $\Delta_{GPi,Str}$ | 8 | $J_{GPi,Str}$ | 1.6 | $\tau_{STN,M_{ctx}}$ | 20 |
| $\Delta_{Tha,GPi}$ | 5 | $J_{Tha,GPi}$ | 0.3 | $\sigma_{GP,STN}$ | 120 |
| $\Delta_{Ctx,Tha}$ | 5 | $J_{M_{ctx},Tha}$ | 1.2 | $\sigma_{M_{ctx},M_{ctx}}$ | 100 |
| $\Delta_{STN,M_{ctx}}$ | 5 | $J_{STN,M_{ctx}}$ | 3.5 | $\sigma_{Str,PPC}$ | 100 |
| $\Delta_{GPi,STN}$ | 5 | $J_{GPi,STN}$ | 3.3 | $\sigma_0$ | 9 |
| $\Delta_{GPi,GPe}$ | 2 | $J_{GPi,GPe}$ | 0.3 | $JE_{M_{ctx},M_{ctx}}$ | 4.5 |
| $\Delta_{GPe,STN}$ | 3 | $J_{GPe,STN}$ | 0.5 | $JI_{M_{ctx},M_{ctx}}$ | 0.6 |
| $\Delta_{STN,GPe}$ | 3 | $J_{STN,GPe}$ | 0.6 | $J_{M_{ctx},PPC}$ | 2 |
| $\Delta_{GPe,Str}$ | 7 | $J_{GPe,Str}$ | 0.5 | $J_{Str,PPC}$ | 2 |

We model each population in the BG-thalamo-cortical network as a network with a ring architecture. Each neuron is characterized by its position on the ring which corresponds to its preferred direction (PD) during reaching movement execution. For simplicity, recurrent connectivity is implemented only in M_ctx_. The strength of the recurrent connections depends on the difference between the pre- and post-synaptic neuron’s PD. In accordance with the topographic organization of the BG-thalamo-cortical network, the same rule also applies to all feedforward projections in the model.

Neurons in the PPC are tuned to the location of visual cues (35). Anatomical and physiological evidence indicates that motor and PPC cortical inputs related to the same movement converge in the striatum (38, 39). We therefore assume in our model that the strength of connections between the PPC neurons and M_ctx_ or striatal neurons depend solely on the difference between the preferred orientation of the pre-synaptic PPC neuron and the post-synaptic neuron’s preferred direction (PD).

We consider reaching tasks where a subject performs arm movements directed by a transient visual cue presented briefly (250 ms). This transient cue evokes a transient directional input in PPC neurons I_cue_(θ_cue_, t) (Fig. 1C). During movement, M_ctx_ neurons also receive a uniform, time-dependent external input, I_mov_(t), reflecting motor drive from premotor areas (Fig. 1C). Finally, we posit that the network’s behavioral output corresponds to reaching movements, with their direction and vigor determined by the activity of the neurons in M_ctx_ (29, 40) (see Material and Methods).

### Circuit activity during a reaching movement

Unless stated otherwise, the model’s parameters are as in Table 1. At rest, I_mov_(t) = 0 and the activity in M_ctx_, the GPi, as well as other populations, is homogeneous (Figs 1D-E). Consistent with experimental observations, the average rest activity is significantly higher in the GPi, GPe, thalamus and STN compared to M_ctx_, while the striatum exhibits lower activity than M_ctx_ (39).

When the cue appears in direction θ_cue_, M_ctx_ and the striatum receive cue-related input encoding this direction from the PPC (Fig. 1C). This input is strongest for striatal and M_ctx_ neurons whose preferred direction is θ = θ_cue_. At movement onset, the input I_mov_(t) gradually increases over 500 ms, peaks, and then decreases over the next 500 ms, forming a bell-shaped temporal profile lasting 1 second after movement initiation (Fig. 1C). During movement, M_ctx_ activity displays a bumpy profile: only a subset of neurons increases their firing rate, while others remain near their spontaneous activity levels (Fig. 1D). This pattern arises despite the uniformity of I_mov_(t) and results from the interplay between recurrent dynamics within M_ctx_, its input from the PPC and the strong feedback from the BG-thalamo-cortical loop. Therefore, the activity in M_ctx_ is biased by a monosynaptic input from the PPC and a polysynaptic input from the striatum through the BG. Note that M_ctx_ consistently points to the cue direction for the whole duration of the movement although I_cue_(*t*) is transient. The selection of the proper movement direction requires I_cue_(*t*) to last >100ms (Supplementary Fig 1).

During the movement, the population vector generated M_ctx_ neuronal activity evolves in time, coding for the trajectory of the hand (see Materials and Methods; Fig. 1F). The movement direction corresponds to the instantaneous direction of the vector. It aligns well with the cue’s direction despite the noise generated by the network dynamics. The movement amplitude is determined by the maximum length of the population vector. The time-derivative of the amplitude corresponds to the velocity of the movement. Movement-related activity reverberates throughout the entire BG-thalamo-cortical loop. As a result, all populations within the BG and the thalamus exhibit heterogeneous, directionally tuned activity patterns (Fig. 1G). The striatum focally inhibits the GPi through the direct pathway, reducing selectively the activity in GPi neurons whose PD aligns with movement direction (Fig. 1E). Conversely, GPi neurons with a PD opposite to movement direction are strongly excited due to the combined effects of divergent excitatory input from the STN and diminished striatal inhibition. This differential activation produces a “bump” of activity in the GPi that is inverted compared to the patterns observed in the cortex, striatum, and STN. Thereby, in our model GPi neurons display sharp tuning curves (Fig. 1G), consistent with experimental observations (41, 42). Inactivation of the GPi does not prevent the movement-related activity in M_ctx_ (Fig 1H), but the peak of activity is reduced, leading to a reduced movement velocity and amplitude, *i*.*e*., reduced movement vigor.

### Cortico-striatal synaptic plasticity drives movement adaptation

We investigated in the model the conditions under which reaching movements can adapt through reward-driven learning. To this end, we considered a hand-reaching task in absence of visual feedback about the actual hand location (43–45). Initially, the “subject” is instructed to move in the direction of a cue, θ_cue_, performing successfully in most trials (Fig. 1). Subsequently, the target location is rotated by an angle Δθ with respect to θ_cue_ (Fig 2A). Importantly, the subject is not informed of this new condition. The only information on performance is a reward provided by the “experimentalist” at the end of a trial, according to the angular distance of the hand with respect to the desired target. Specifically, the rewarded interval is [θ_cue_ − Δθ, θ_cue_ + Δθ]. At the end of the n^th^ trial, the weight of the cortico-striatal projection from a PPC neuron with PD, θ_*PPC*_, and a striatal neuron with PD, θ_str_, changes according to the reward-modulated tri-partite learning rule (46, 47) (see Material and Methods for details)

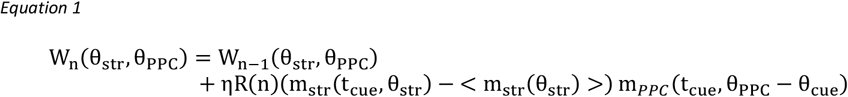

where η is the learning rate, R(n) the value (0 or 1) of the reward at the end of the trial. Here, m_str_(t_cue_, θ_str_) and m_PPC_(t_cue_, θ_PPC_ − θ_cue_) denote the activity of the striatal and PPC neurons when the cue appears in the n^th^ trial in the direction θ_cue_. Finally, < m_str_(θ_str_) > is the time average of the striatal neuron activity over the 500 msec preceding the onset of the cue.

**Figure 2:**
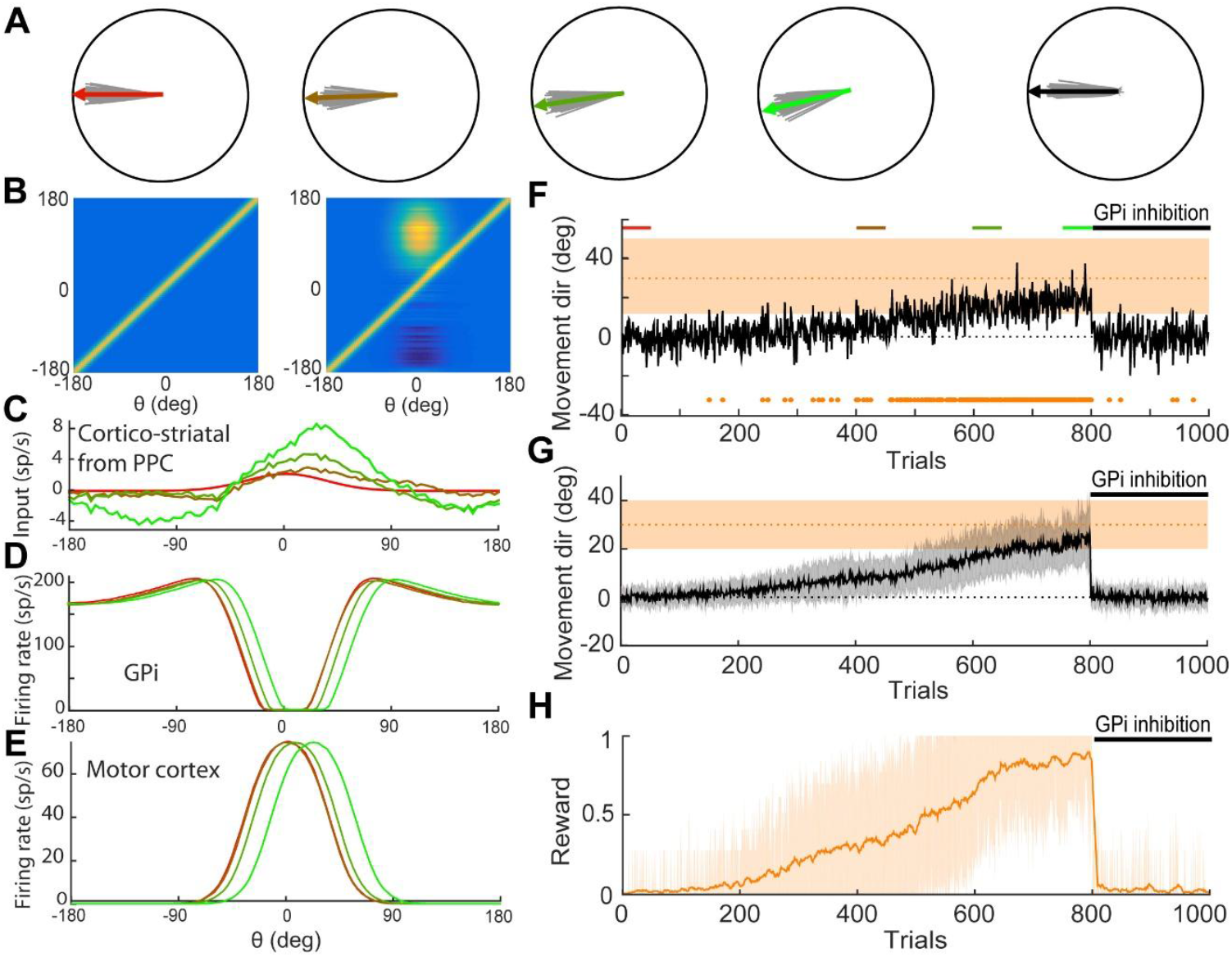
Reward-modulated cortico-striatal plasticity allows adaptation of movement direction in a reward-driven rotation protocol with 30° shift. GPi inhibition after learning cancels the learned adaptation bias. A: direction of movements produced along the reward-driven adaptation protocol. Example trajectories of 50 movements toward the target at different stages of the adaptation protocol are shown in gray, with mean movement trajectory in color (from left to right: early, red, trials 1-50; intermediate, brown, 401-450; intermediate, dark green, 601-650; final, light green, 751-800; after GPi inhibition, black, 800-1000). B: Matrix of synaptic weights between M_ctx_ and the Striatum before (left) and after(right) adaptation. C-E: Profile of the input from the PPC to striatal neurons (C) and activity of the GPi neurons (D) and cortical neurons (E) across trials along the adaptation protocol. Example trials are coloured according to their position in the protocol (red: early, green: late). F: Movement direction along the adaptation protocol in an example run. The orange shaded area indicates rewarded movement directions. Orange points along the x axis indicate rewarded trials. In trials 800-1000 the GPi is inhibited (100% inhibition). G-H: Average movement direction (G) and reward rate (H) over 20 runs of the adaptation protocol. The orange shaded area in G indicates rewarded movement directions. The gray shaded areas indicate the mean +/- SD of the movement direction in G. The orange shaded area indicates the mean +/- SD of the reward rate in H.

Prior to adaptation, the combined excitatory input that striatal neurons receive from the PPC is biased in the direction of the cue. This bias arises because the projections from the PPC to the striatum are directionally specific (Fig 2B, left, and red line in Fig 2C). During adaptation, these projections are adjusted in accordance with Equation (1). Consequently, their functional specificity shifts, and the directional bias of the aggregate input from the PPC to the striatum realigns toward the rewarded desired direction (Fig. 2C).

Figures 2D-F, illustrate how the adaptation is driven by the BG-cortical network. Due to the inherent noise in movement production, the hand explores a range of final reach positions. Movement direction occasionally falls within the rewarded interval, leading to positive reinforcement. The projections from neurons in the PPC to the striatum that are selective for the rewarded directions are strengthened. Because the projections along the direct pathway are functionally specific, neurons in the GPi, thalamus, and M_ctx_ coding for the rewarded direction undergo inhibition, disinhibition, and activation, respectively. This is demonstrated for GPi and M_ctx_ neurons in Fig. 2D&E (green lines). As a result, the aggregate polysynaptic input from the BG-thalamic circuit to M_ctx_ becomes increasingly biased as learning progresses. Consequently, the likelihood of the hand movement ending within the rewarded interval increases, leading to a higher fraction of rewarded trials (Fig. 2G&H).

The averaged learning curve over twenty repetitions of the protocol (Fig. 2G&H) shows that by the end of learning, the movement has successfully adapted to the new condition. This adaptation critically depends on the bias in the input from the BG-thalamic circuit to M_ctx_ toward the desired movement direction. Suppressing this bias by inhibiting the GPi disrupts the adaptation, resulting in its loss (Fig. 2F-H). As mentioned above, GPi inhibition also reduces the vigor of movement and thus the effect of the synaptic noise on movement direction is magnified, leading to a slightly more variable movement direction following GPi inhibition.

### Strong cortico-BG motor loops support efficient adaptation

As just described, during adaptation, changes in the input from the PPC to striatal neurons modify the movement-related activity patterns in the GPi. This, in turn, causes the feedback input from the striatum to M_ctx_ to bias the population vector generated by M_ctx_ neurons toward the desired movement direction. How does this depend on the strength of the projections in the direct loop? How does the efficiency of the learning process vary with these parameters?

To address these questions, we simulated the model with varying strengths of cortico-striatal projections. The results show that as the strength of these projections decreases, the learning dynamics slow down (Fig. 3A). Moreover, the average rotation angle of the movement direction becomes smaller (Fig. 3B). When the cortico-striatal projections are too weak, the probability of success during learning drops drastically and the movement direction shows minimal change by the end of a block of trials. These findings highlight the importance of the feedback dynamics in the BG-thalamo-cortical loop for effective movement adaptation.

**Figure 3:**
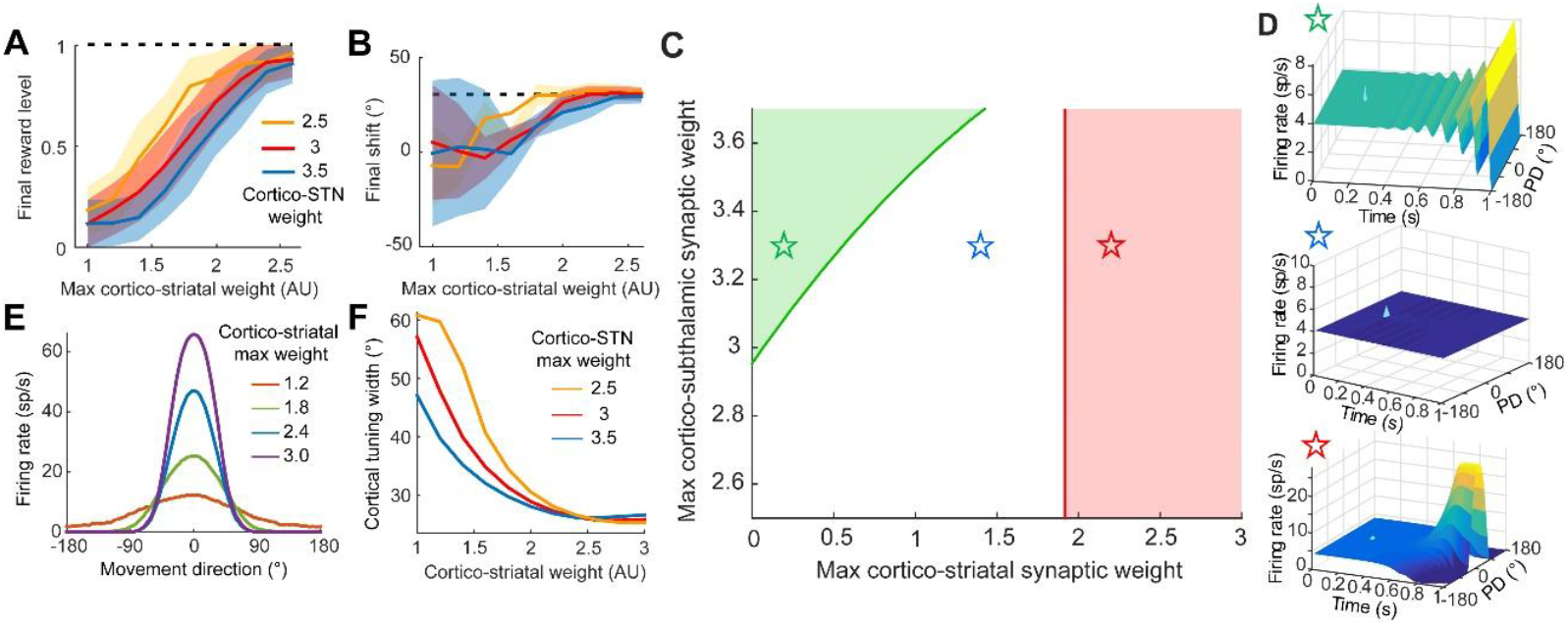
Cortical movement-related activity profile and the network ability to adapt movement direction strongly depends on the average cortico-striatal synaptic weight. A-B: The success of movement adaptation (target at 30° +/- 20°) is compromised when the cortico-striatal synapses are weakened, as evidenced by final reward level (A) or final movement direction after 1000 trials (B). This is true for various cortico-STN maximal synaptic weights (different colors as indicated). C: Phase diagram showing the three dynamical regimes of the BG-cortical network as a function of the maximal cortico-striatal (G_str,Mctx_) and cortico-subthalamic (G_STN,Mctx_) synaptic weights. For each value of (G_str,Mctx_, G_STN,Mctx_), the network is set at the steady state of its dynamics, where constant external inputs lead to constant activity levels in all populations. A perturbation is applied to this steady state to test its stability, highlighting the properties of the possible instabilities of the network. The white area corresponds to the domain where the steady state is stable and the network remains at constant activity levels after perturbation. The red line indicates a symmetry breaking instability while the green line denotes an oscillatory instability. Both lines are computed analytically from the equations governing the network stability (See Material&Methods). D: Cortical activity profile following the perturbation of the network from its steady state. Top: oscillatory instability leads to cortical oscillations that propagate through the network. Middle: Stable steady state. Bottom: symmetry breaking instability leads to a sharp activity profile in the cortical population due to the network dynamics. E: Cortical tuning curve is narrow for strong cortico-striatal weights and broad when cortico-striatal connection is weakened. F: The decrease in cortical tuning with the maximal cortico-striatal weight is true for a large range of cortico-STN synaptic weights.

To further explore the role of the BG-cortical loop in the adaptation process, we analyzed the network dynamics in the absence of recurrent interactions within M_ctx_. All parameters were fixed except for the M_ctx_-striatal synaptic weights (Fig. 3C-D). Under these conditions, the BG-cortical loop exhibit three distinct dynamical regimes. For intermediate values of the cortico-striatal weights, the network remained in a stable homogeneous activity state. In this regime, activity in M_ctx_ is uniform (Fig. 3D, middle). As the cortico-striatal weights increase, this homogeneous state losses stability. A symmetry-breaking bifurcation occurs (Fig. 3D, bottom). This results in heterogeneous ‘bump’ activity profile within M_ctx_. Notably, this bifurcation barely depends of the negative feedback mediated by the hyperdirect and indirect pathways (Fig. 3C). This is because the projections from STN to the GPe and GPi are broadly tuned (Material&Methods). For sufficiently small cortico-striatal synaptic weight, a bifurcation occurs to a state in which the activity of neurons synchronously oscillates across all populations. These oscillations are driven by the hyperdirect negative feedback loop (11). This analysis underscores how the cortico-striatal synaptic weights shape the dynamics of the BG-cortical loop.

We then examined how the recurrent dynamics in the M_ctx_ network interact with the BG-thalamo-cortical dynamics to shape movement-related activity in M_ctx_. Figure 3E illustrates how the movement-related activity profile in M_ctx_ changes with the strength of the M_ctx_-striatal projections. When these projections are weak, the activity in M_ctx_ is broadly tuned. It sharpens as these projections strengthen. As explained above, the intrinsic dynamics of the BG-cortical loop undergo a bifurcation to a bump state for sufficiently strong M_ctx_-striatal projections, further sharpening M_ctx_ activity profile and amplifying its activity. This effect is robust to variations in other model parameters, as shown in Fig. 3F for changes in cortico-subthalamic synaptic weights.

In summary, when M_ctx_-striatal projections are weak, the influence of BG-thalamo-cortical feedback on M_ctx_ is minimal, and the bias toward the cue induced by *PPC*-M_ctx_ projections dominates (Fig. 3B). Conversely, when M_ctx_-striatal projections are strong, the bias induced by BG input dominates and shifts the activity bump in M_ctx_. Since during learning this bias rotates toward the desired direction, the population vector in M_ctx_ progressively aligns with the target direction (Fig. 3A-B).

### Shaping adaptation in the low-noise regime

In our model, the synaptic noise drives exploration in movement trajectories. Thus, the learning rate strongly depends on the noise level (Fig. 4A-D). In particular the level of noise determines the time to achieve the first reward. At very low noise levels (orange), learning is extremely slow, with fewer than 1% of movements reaching the reward region after 1,000 trials. For larger noise (red), the initial learning phase is shorter, enabling faster attainment of the first reward. Subsequently, adaptation accelerates, achieving an average final performance above 80%. However, further increasing the noise (blue) speeds up initial learning but reduces final performance due to imprecise movements. Excessive noise disrupts movement accuracy both before and after adaptation, with its amplitude remaining stable throughout reward-driven learning(48, 49). This trade-off between sufficient exploration and movement precision constrains adaptation efficiency.

**Figure 4.**
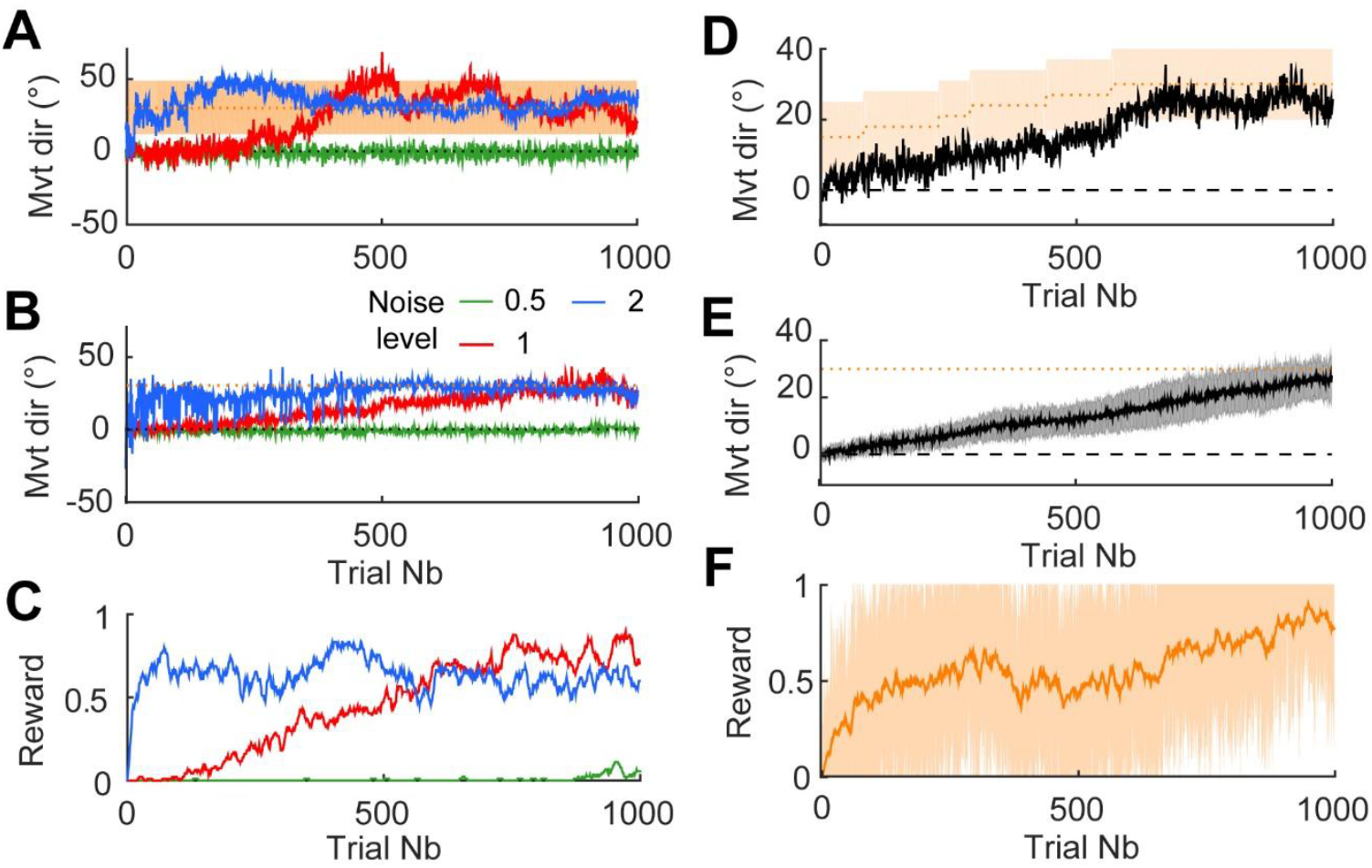
Adaptation is very slow for low movement variability, and adaptation in realistic conditions requires shaping. A: Example learning curves for adaptation of movement direction in the model with various levels of striatal synaptic noise (0.5, 1 and 4). The shaded area represents rewarded movement directions (30° +/- 20°). While low noise level lead to low movement variability and thus prevent adaptation, high noise leads to fast convergence. B-C: Mean learning curves (B) and reward ratios (C) for striatal synaptic noise levels of 0.5-1-2. D: Example learning curve during an adaptation protocol including shaping. The shaded area represents rewarded movement directions. The direction of the target (and thus rewarded movements) is slowly increased as the movement direction is modified. E-F: Mean learning curves (E) and reward ratios (F) over 20 runs in the shaping protocol. The shaded area indicates the mean +/- SD of the reward rate.

Behavioral experiments show that baseline reaching movements are typically accurate (43, 44), indicating low noise levels and slow adaptation. A “shaping” protocol can mitigate this limitation and enhance adaptation (45, 50) (Fig. 4D-F). To accelerate learning, one strategy would be to progressively increase the noise level, though implementing this experimentally is challenging. An alternative approach is to shape behavior by gradually increasing task difficulty (50). In such a protocol, the target direction is adjusted dynamically based on current performance, as shown in Fig. 4E-H. Initially, the rewarded area is close to the cue direction (15° ± 10°), resulting in frequent rewards. Reward-driven cortico-striatal plasticity then shifts movements progressively toward the desired target. As the target direction is gradually updated, movements adapt continuously toward the final goal (Fig. 4E-F). This method enables faster adaptation, even under low noise conditions, without compromising the accuracy of baseline reaching movements (Fig. 4G-H).

### Basal ganglia drive *de-novo* visuo-motor association

The association between visual cues and movement direction is not innate (51); rather, visuo-motor associations are acquired during early childhood. More generally, sensorimotor associations develop through spontaneous exploration of movement possibilities, a process commonly referred to as “babbling” (51, 52). We used our model to examine the hypothesis that the basal ganglia (BG) play also a critical role in the initial acquisition of visuo-motor adaptations (53). To this end, we considered a “tabula rasa” BG-thalamo-cortical network, where initial projections from the PPC to the striatum are non-specific (Fig. 5B, red), and projections from the PPC to M_ctx_ are absent. In this configuration, the representation of cue and movement directions in the network are independent. We nonetheless assumed that neurons in the PPC are tuned to the location of visual cues and that projections from M_ctx_to the striatum are topographically organized. As illustrated in Fig. 5, the alignment between cue and movement direction emerges through the interaction of reward-driven cortico-striatal synaptic plasticity and the dynamics of BG-thalamo-cortical circuit (See Materials & Methods).

**Figure 5.**
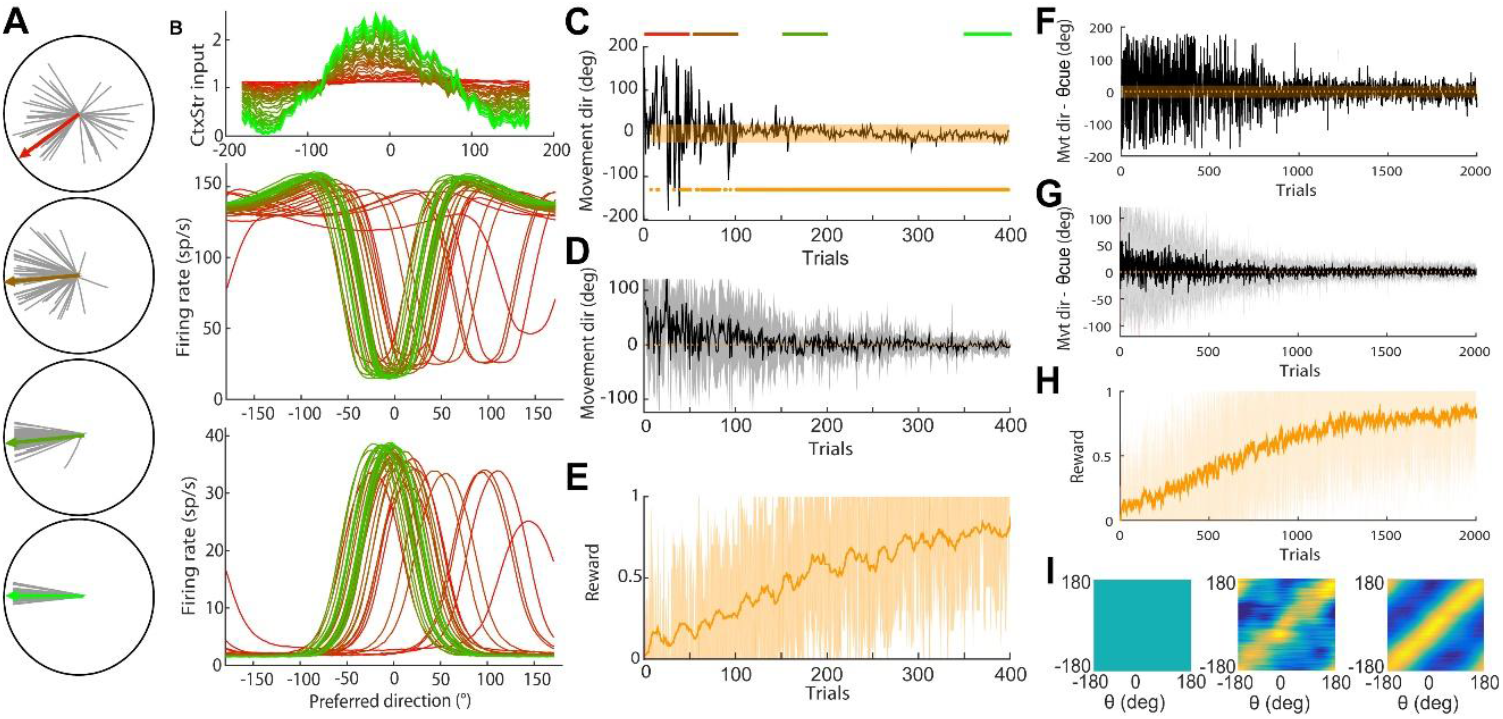
De novo learning: aligning visual cue direction with movement direction of spontaneous reaching movements in a system with no cortico-cortical connectivity. A: direction of movements produced along the reward-driven adaptation protocol. Example trajectories of 20 movements toward the target at different stages of the initial learning protocol (from left to right: trials 1-20, 101-120, 201-220, 381-400). B: Profile of the input from PPC to the striatal neurons (top panel) and activity of the GPi neurons (middle panel) and cortical neurons (bottom panel) across trials along the initial learning protocol. Example trials are colored according to their position in the protocol (red: early, green: late). C: Movement direction along the adaptation protocol in an example run. The shaded area indicates rewarded movement directions. Points along the x axis indicate rewarded trials. D-E: Average movement direction (D) and reward rate (E) over 20 runs of the initial learning protocol. Shaded areas indicate the mean +/- SD of the movement direction (in D) or reward rate (in E). F-I: De novo learning of 16 directions, with multiple cues presented in a random order and uniformly distributed between 0 and 360 degrees. F: During the learning process, movements directed toward the presented cue are rewarded, and movements progressively align to cue directions. G-H: For 20 runs of the learning of 16 direction, the learning is efficient in around 1000 trials, as shown with movement direction with respect to cue direction (G) and reward rate (H). I: During the process, the synaptic weights from PPC to striatum, initially non-specific (left) become structured (final matrix of PPC-striatum synaptic weights after 200 trials: example run, middle; average over 20 runs, right).

We first considered a scenario in which the same cue is repeatedly presented in a fixed direction. Initially, projections from the PPC to the striatum are non-specific, meaning that the sensory input reaching striatal neurons during movement (as described in Fig. 1) does not favor any direction. Meanwhile, the dynamics of BG-thalamo-cortical circuit generate a localized activity bump in M_ctx_, with its position dictated by ongoing synaptic noise. Consequently, movement direction varies randomly across trials, exploring many directions (Fig. 5A, top). When movements align with the presented cue direction (θ_cue_) and fall within a rewarded range θ_cue_ ± 20° (shaded area in Fig. 5C), the projections from the PPC neurons selective to the cue location to striatal neurons tuned to that movement direction become gradually stronger. This leads to the development of direction-specificity in the projections from PPC to striatum (Fig. 5B, top). Due to the topographic organization of striato-pallidal projections, neurons in GPi that are preferentially active during movement in direction θ_cue_ are inhibited. Since the cortico-BG-thalamo-cortical loops are closed, cortical neurons tuned to θ_cue_ increase their activity, while less-tuned neurons are suppressed. As a result, movement direction variability decreases (Fig. 5C&D), making it more likely that future movements fall within the rewarded range (Fig. 5E). Repeating this process over multiple trials further strengthens the projections from PPC to striatum in a direction-specific manner (Fig. 5B).

Next, we considered a case where multiple cues are presented randomly in 16 possible directions evenly spaced between 0 and 360 degrees (Fig. 5F-H). Initially, projections from the PPC to the striatum are non-specific, resulting in random movement directions that are independent of the cue. During learning, movements that align with the cue direction are rewarded. As a result, the projections from the PPC to the striatum are selectively strengthened according to Equation (1). Over time, the connection matrix from PPC to striatum becomes more structured, gradually promoting movements that align with the cue (Fig. 5F). After 2000 trials, movements fall within θ_cue_± 5° in most cases (Fig. 5F), leading to an average reward rate exceeding 80% (Fig. 5G-H). The structure of the final synaptic weight matrix between the PPC and the striatum reflects a strong alignment between the pre-synaptic cue direction in the PPC and the post-synaptic movement direction in the striatum (Fig. 5I).

## Discussion

We have developed a simplified dynamical model of the BG-thalamo-cortical network to investigate the role of BG in reward-based adaptation of reaching movement direction. The model is consistent with the fact that inhibiting the output of the BG impacts the vigor of well-learned reaching movements without affecting their direction. A key result of the model is that the ability of the network to drive adaptation depends on the dynamical regime in which the BG network operates. Our model predicts that after adaptation, inactivation of the output nucleus of the BG (the GPi), lead to a reversion to baseline performance.

### Error-based *vs* reward-driven sensorimotor adaptation

Error-based adaptation relies on sensory feedback to correct predictable perturbations on a trial-by-trial basis (54–56). It is likely mediated by cerebellar circuits (57, 58). When sensory feedback is available, error-based corrections tend to dominate the adaptation of reaching movements (59, 60). In contrast, reward-driven adaptation relies on a binary evaluation of action outcomes. This form of adaptation has been studied in humans performing motor tasks where outcomes are rewarded based on rules known only to the experimenter. Reward-driven sensorimotor adaptation can pertain to various movement features, such as curvature (48) or direction (44, 45). This form of adaptation remains intact in individuals with cerebellar damage (45) but requires functioning BG circuitry (1, 4, 17, 18). Furthermore, dopaminergic input to the BG is both necessary and sufficient for facilitating rapid reinforcement-driven behavioral corrections (5, 61, 62). The mechanisms by which BG circuitry and its dynamics contribute to reward-based adaptation are still largely unclear. Our work seeks to address this issue using a biologically-inspired model of the cortex-BG network. It demonstrates how the representation of reaching movements in the BG can align with the target direction after adaptation. The underlying adaptation relies on the combination of a tri-partite cortico-striatal Hebbian plasticity modulated by the reward signal at the PPC-striatal synapses with the feedback dynamics of the loops embedded in this network.

### Comparison with other computational models

Standard reinforcement learning (RL) algorithms are effective for learning discrete tasks (e.g., choosing between two options)(63). Implementation of RL relying in the dorsal BG circuits have been proposed in previous modelling work (3, 64). In these models, actions are represented by discrete populations in BG output nuclei (3, 65–68). In the present work, we have developed a mechanistic model for the role of BG in learning simple continuous movements.

In current theoretical models of birdsong RL (69, 70) the reward-modulated plasticity occurs at cortico-cortical synapses. However, it has been found that dopamine-dependent plasticity primarily takes place in cortico-striatal connections which subsequently drive BG-dependent motor corrections within cortical networks (17, 71). Tesileanu et al. (72) proposed a mechanism whereby a cortico-cortical network learns from another cortical area acting as a “tutor,” which has been trained through reinforcement learning. This work is complementary our study since it addresses how cortico-cortical connections slowly learn adaptations initially implemented through reward-driven by BG output.

The mechanism we propose for the role of the BG-cortical network in reward-based sensorimotor adaptation differs fundamentally from previous theories (3, 71). It is based on the emergence of attractors within the recurrent dynamics of the BG-cortex circuit. An integration of reward-driven learning and attractor dynamics has been previously proposed in the context of operant conditioning, where subjects choose from discrete options (73). In our model, these attractors arise from feedback loops that topographically connect the cortex to itself via the basal ganglia and thalamus, as shown in anatomical studies (22–24) and more recent physiological experiments (22). These experiments revealed that specific subsets of cortical information are transmitted through the cortico-BG -thalamic network back to the original cortico-striatal neurons of each subnetwork, thereby forming a bona fide closed loop (22).

In our model, reaching movements are represented by the population vector of neuronal activity in the motor cortex (64). The vigor and direction of the movement correspond to the norm and direction of this vector. A different view of movement representation in the motor cortex, known as the “dynamical system perspective” of movement production, posits that the geometry of the trajectory in the multidimensional space spanned by single neural activity space determines the muscle activation patterns (77, 78). Several computational models of movement production have been developed according to this view(79, 80). They explain the changes in the direction of the vector representing motor cortex activity in neural space from preparation to execution of a movement (81). However, such changes can also be accounted for in models with functional organization (40). Moreover, a careful analysis of the response to perturbation in the activity of motor cortex neurons shows that they encode reaching movement direction through a low-dimensional attractor (82).

Our model does not consider the opposing role of the direct and indirect pathway in the BG during action selection and learning (83). Interestingly, a recently proposed mechanism implementing an opposing learning rule in striatal neurons engaged in each of these two pathways (84) could be compatible and complementary to the network mechanisms highlighted in this study. We intentionally omit recurrent connectivity in the BG nuclei for the sake of simplicity of the model. While recurrent connectivity is relatively weak in the STN and thalamus, it is more pronounced in other BG nuclei, such as the GPe (21). Understanding the functional implications of this recurrent connectivity would necessitate further theoretical investigation.

### Generality of the results

While our model is grounded in the anatomy of the BG and reaching-related activity in primates(29), the principles it reveals are general and may broadly apply to BG-cortical circuits in vertebrates. These principles rely on two key features: (i) topographically organized closed-loop BG-cortical circuits, which have been observed in rodents, songbirds, and non-human primates (NHPs)(22–24); and (ii) pathways of opposing polarity within the BG, a feature present in species as diverse as lamprey, songbirds, rodents, and NHPs, with a remarkably well-conserved functional connectivity (37, 85–87). This suggests that similar network dynamics may underpin movement production and adaptation across a variety of vertebrates. The relative contribution of the BG and their target action generation circuits in the cortex or brainstem (8, 9, 28) may however differ in the various species and sensorimotor modalities. As an example, saccadic eye movement remain BG-dependent in adult monkeys (88). The BG-driven motor learning mechanism proposed may be apply in these various BG circuits as long as the network is organized in parallel feedback loops.

In our model, the topographic projections to the motor cortex from a population of neurons selective to the direction of the visual cue play a crucial role in aligning the sensory and motor representations of movement direction. We demonstrate that this alignment can emerge de novo through the same plasticity rules when paired with large-scale exploratory behavior, such as motor babbling (Fig. 5), similar to vocal babbling observed in infants and songbirds (51, 52). In this view, babbling allows young subjects to link high sensory representation of a goal to the motor command required to reach it during early motor development(89, 90).

### Predictions and perspectives

Like all models of reward-based sensorimotor adaptation involving the BG, we predict that inhibiting the GPi will disrupt adaptation. In our framework, reward-driven adaptation induces synaptic changes in cortico-striatal projections but does not alter recurrent connectivity in the motor cortex. Therefore, we predict that if the output of the BG to the motor cortex is suppressed shortly after learning, performance will revert to baseline. This could be tested in NHPs trained without visual feedback, guided only by external rewards (*e*.*g*., as in (91)), by pharmacologically or chemogenetically inhibiting the GPi (as shown in Fig 2C). A second prediction of our model is that adaptation will also be impaired by weakening the projections from the motor cortex to the striatum. This could be tested in NHPs by expressing chemogenetic receptors in motor cortex projection neurons (92), and locally activating these receptors in the dorsal striatum. This would reduce synaptic release by cortico-striatal axons in the striatum without affecting other movement control centers. Our theory can also be tested in the context of pitch adaptation in songbirds (17). Here, chemogenetic methods could be used to reduce the strength of projections from the Lateral Magnocellular Anterior Nucleus, a cortical region, to Area X, the songbird equivalent of the BG (striatum, globus pallidus). Our model predicts that this reduction would impair pitch adaptation.

## Materials and Methods

### Architecture of the model

Our model comprises seven neuronal populations (Fig. 1A): the striatum, the internal and external segments of the globus pallidus (GPi and GPe), the subthalamic nucleus (STN), the thalamus and two cortical populations. One cortical population represents M1 (M_ctx_) and the other one, PPC, represents to the posterior parietal cortex. In each population, neurons are distributed on a ring. The location of a neuron is its preferred direction (PD) during reaching movement execution (40, 74, 93), denoted as an angular variable θ, except in the PPC neurons, where the PD of a PPC neuron is defined as the direction of the cue for which its response is maximum. The strength of the connections between pre- and post-synaptic neurons from two interconnected neuronal populations depends on the difference between their PD. Specifically, the connection strength between neuron (*θ, α*) and (*θ*^′^, *β*) is

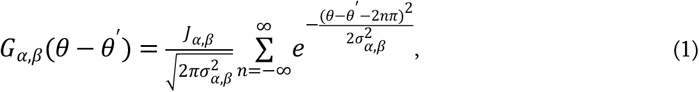

where α, β ∈ (Mctx, str,GPi,GPe,STN,tha, PPC) are the populations the pre and the post synaptic neuron belong to. Here σ_α,β_ is the dispersion (footprint) of the connections between the two populations. Note that in accordance with the ring architecture of each population the function *G*_α,β_(Δθ) is period (period 2π). The normalization factor is such that its integral is J_α,β_. For the sake of simplicity recurrent connectivity are present only in M_ctx_. Given the tight topography of connectivity along the direct pathway and its thalamo-cortical feedback (22–24), we consider all dispersion factors along these connections to be the same (σ_α,β_ = σ_0_ for (α, β) ∈ ((str,Mctx), (GPi,str), (GPe,str), (GPi,GPe), (tha,GPi), (Mctx,thal), (STN,Mctx))) and low compared to other connections (see Table 1 for parameter values).

### Neuronal dynamics

Neurons are modeled as rate units (94, 95) in which the dynamical variables describe the synapses (95). Denoting by I_α_(θ, t) the total input into the neuron in population α and PD, θ at time t, its instantaneous firing rate, A_α_(θ, t), in spikes/s, is

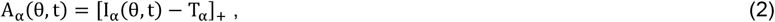

with [x]_+_ = x for x > 0 and 0 otherwise and T_α_ is a threshold which depends only on the population.

The synaptic current induced by neuron (θ′, β) into neuron (θ, α) is *G*_α,β_(θ − θ^′^)mαβ(θ, t) where the synaptic activity, m_αβ_(θ, t), satisfies (95, 96)

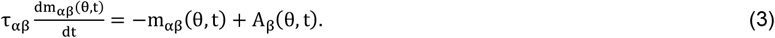

Here τ_αβ_ is the time constant of the synapses that neurons in population β make on neurons in population α . For simplicity, we assume that τ_αβ_ does not depend on *α* and β, τ_αβ_ = τ, but for (α, β) = (STN, Ctx) for which the synaptic time constant is larger to account for the presence of NMDA receptors on STN neurons. As a result, m_αβ_(θ, t) does not depend on *α* unless (α, β) = (STN, Ctx). But in the latter case, we will write m_αβ_(θ, t) ≡ m_β_(θ, t) to simplify the notations.

### Inputs to the neurons

All the populations receive a movement independent constant external input, *H*_α_ as well as well as a Gaussian white noise, η_α_(θ, *t*), with zero mean and SD, *s*_α_. This noise is uncorrelated across neurons. The striatal population receives an additional input noise, *ξ*_*str*_ (*θ, t*), which is correlated in time and across neurons. This noise which drives the exploration during learning is determined by

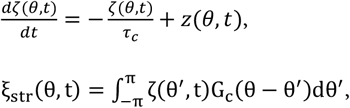

where z(θ, t) is a Gaussian white noise with SD, *s*_c_, which is uncorrelated across neurons, τ_*c*_ is the correlation time and

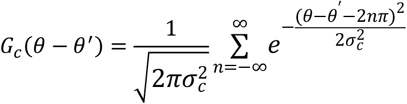

We consider reaching a task in a direction, instructed by a transient visual cue. This cue evokes a transient directional selective sensory input, I_cue_(θ_cue_ − θ, t) into a PPC neurons of preferred direction θ. Here, θ_cue_ is the direction of the cue. During movement, M_ctx_ neurons also receive a uniform, time-dependent external input, I_mov_(t) corresponding to a motor drive from premotor areas. It does not depend on the movement direction.

Considering the architecture of the model the total inputs into the neurons read

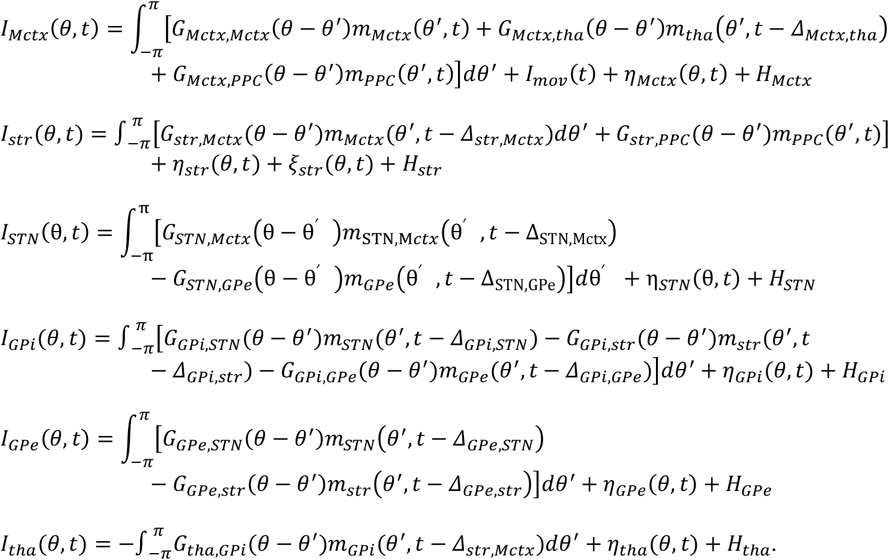

where Δ_α,β_ is the synaptic delay from population *α* to population *β*.

### Movement execution

Neuron activation in the motor cortex directly drives movement. In our model, the instantaneous movement direction and speed are determined by a population vector. This vector is calculated as the sum of individual neurons’ vectors, each pointing toward its preferred direction (29, 40), with the amplitude proportional to its activity above 10 spikes per second (sp/s). The 10 sp/s threshold ensures no movement occurs during typical cortical rest activity, which averages around 5 sp/s. A movement is triggered as soon as any neuron exceeds the 10 sp/s activity level. We assume that during the planning and execution of movements, neurons in both cortical and striatal populations receive additional external inputs. These inputs represent synaptic signals arriving at cortical and striatal neurons from other cortical areas and brain regions that are not explicitly included in the model. For simplicity, the inputs to the cortical populations are considered homogeneous across the entire population. In both the text and figures, external inputs are expressed in arbitrary units.

### Integration method

The model’s dynamics were simulated using a standard first-order Euler algorithm with a fixed time step of 0.5 ms. To ensure the accuracy of the simulations, additional runs were performed using a smaller time step (0.05 ms), confirming that the dynamics remained consistent and unchanged.

## Acknowledgments

AL thanks Alexis Dubreuil, Nicolas Mallet and Marc Deffains for thoughtful discussions. DH expresses gratitude to Carole Levenes, Yonatan Loewenstein, and the late Carl van Vreeswijk for their insightful discussions. He also thanks the Edmond and Lily Safra Brain Research Center, and the Loewenstein’s lab at The Hebrew University of Jerusalem, Israel for their warm hospitality. This work was supported by funding from the Agence National pour la Recherche (16-CE37-0020-01, 20-CE37-0023-02, 22-NEUC-0002-01), Fonds pour la Recherche Médicale (EQU202203014688) to AL, and the France Israel Center for Neural Computation (FICNC, HUJI/CNRS) to DH.

