## Supplementary methods and figures for "Reward-driven adaptation of movements requires strong recurrent basal ganglia-cortical loops"

#### 1 Derivation of the bifurcation lines in Fig. 3C

We outline here the derivation of the instability (bifurcation) lines of the homogeneous fixed-point state of the model in the case of constant external input (uniform in space and time) and in the absence of external noise (Fig. 3C).

The dynamical equations read (see Methods, Eqs. (1–3)):

$$\tau_{\beta\alpha}\,\frac{dm_{\beta\alpha}}{dt}(\theta,t) = -m_{\beta\alpha}(\theta,t) + A_\alpha(\theta,t)\,, \tag{1}$$

where $\alpha,\beta \in \{\mathrm{ctx},\mathrm{stn},\mathrm{str},\mathrm{th},\mathrm{gpi},\mathrm{gpe}\}$.

$$A_\alpha(\theta,t) = \big[I_\alpha(\theta,t) - T_\alpha\big]_+$$

with

$$I_\alpha(\theta,t) = \sum_\beta \int_{-\pi}^{\pi} J_{\alpha\beta}\, G(\theta-\theta',\sigma_{\alpha\beta})\, m_{\alpha\beta}(\theta',t-\Delta_{\alpha\beta})\, d\theta' + H_\alpha\,. \tag{2}$$

The sum runs over all presynaptic populations $\beta$, and the connectivity kernel is defined as

$$G(\theta,\sigma) = \frac{1}{\sqrt{2\pi\sigma^2}} \sum_{n=-\infty}^{\infty} \exp\!\left(-\frac{(\theta-2n\pi)^2}{2\sigma^2}\right). \tag{3}$$

This system always admits a homogeneous fixed-point solution such that, for all $\alpha$ and $\beta$,

$$m_{\beta\alpha}(\theta,t) = m^{\mathrm{FP}}_{\beta\alpha}\,,$$

does not depend on $\theta$ and $t$. However, depending on the network parameters, this fixed point can become unstable. To determine the values of the parameters for which this happens, we study the dynamics of small perturbations around the fixed point state [1]. Stability requires that such perturbations decay with time.

We therefore write

$$m_{\beta\alpha}(\theta,t) = m_{\beta\alpha}^{\mathrm{FP}} + \delta m_{\beta\alpha}(\theta,t)\,, \tag{4}$$

with

$$\delta m_{\beta\alpha}(\theta,t) \ll m_{\beta\alpha}^{\mathrm{FP}}\,.$$

#### Linearization of the dynamics

Substituting Eq. (4) into Eq. (1) and keeping only first-order terms in the perturbations, we obtain

$$\tau_{\beta\alpha}\,\frac{d}{dt}\,\delta m_{\beta\alpha}(\theta,t) = -\,\delta m_{\beta\alpha}(\theta,t) \;+\; \delta I_{\alpha}(\theta,t)\,, \tag{5}$$

where

$$\delta I_{\alpha}(\theta,t) = \sum_{\beta'} \int_{-\pi}^{\pi} J_{\alpha\beta'}\, G(\theta-\theta',\sigma_{\alpha\beta'})\, \delta m_{\alpha\beta'}(\theta',t-\Delta_{\alpha\beta'})\, d\theta'\,, \tag{6}$$

#### Fourier mode decomposition

The function $G(\theta,\sigma)$ is $2\pi$-periodic in $\theta$. So is the case for $m_{\beta\alpha}(\theta,t)$. Writing its Fourier expansion

$$\delta m_{\beta\alpha}(\theta,t) \;=\; \sum_{k\in\mathbb{Z}} \hat{m}_{\beta\alpha}(k,t)\, e^{ik\theta}\,. \tag{7}$$

Eqs. (5)–(6) yields, mode by mode,

$$\tau_{\beta\alpha}\,\frac{d}{dt}\,\hat{m}_{\beta\alpha}(k,t) = -\,\hat{m}_{\beta\alpha}(k,t) \;+\; J_{\alpha\beta'}\,\hat{G}(k,\sigma_{\alpha\beta'})\;\hat{m}_{\alpha\beta'}(k,t-\Delta_{\alpha\beta'})\,, \tag{8}$$

where

$$\hat{G}(k,\sigma) \;=\; \exp\!\left(-\tfrac{1}{2}k^2\sigma^2\right).$$

Seeking exponential solutions $\hat{m}_{\beta\alpha}(k,t) = u_{\beta\alpha}(k)\, e^{\lambda t}$, where $\lambda$ is a complex number Eq. (8) becomes

$$\left(1+\lambda\,\tau_{\beta\alpha}\right) u_{\beta\alpha}(k) \;=\; \sum_{\beta'} J_{\alpha\beta'}\,\hat{G}(k,\sigma_{\alpha\beta'})\; e^{-\lambda\Delta_{\alpha\beta'}}\, u_{\alpha\beta'}(k)\,. \tag{9}$$

For simplicity, we assume that $\tau_{\beta\alpha}$ is independent of $\beta$ except when $\alpha = \mathrm{ctx}$ (Table 1). As a result, $u_{\beta\alpha}(k)$ does not depend on $\beta$ unless $\alpha = \mathrm{ctx}$. We therefore set $u_{\alpha}(k) \equiv u_{\beta\alpha}(k)$ for $\alpha \neq \mathrm{ctx}$. Denoting by $\mathbf{U}(k)$ the seven-dimensional vector whose components are $u_{\mathrm{str\,ctx}}(k)$, $u_{\mathrm{stn\,ctx}}(k)$, $u_{\mathrm{stn}}(k)$, $u_{\mathrm{str}}(k)$, $u_{\mathrm{th}}(k)$, $u_{\mathrm{gpi}}(k)$, and $u_{\mathrm{gpe}}(k)$, the linear system, Eq. (9), can be written as:

$$\mathbf{Q}(k,\lambda)\,\mathbf{U}(k) = 0 \tag{10}$$

For given $k$, Eq. (10) has a non-trivial solution ($\mathbf{U}(k) \neq 0$) only when $\lambda$ is a solution of the transcendental equation

$$D(k, \lambda) = 0 \tag{11}$$

where $D(k, \lambda) = \det \mathbf{Q}(k, \lambda)$ depends on the parameters of the model.

In general, the solutions of Eq. (11) are complex. Loss of stability occurs when, for some $k$, there exists a solution with $\mathrm{Re}(\lambda) = 0$. Substituting $\lambda = i\mu$ into Eq. (11) and separating real and imaginary parts yields two equations parameterized by $\mu$:

$$\mathrm{Re}\, D(k, \mu) = 0 \tag{12}$$

$$\mathrm{Im}\, D(k, \mu) = 0 \tag{13}$$

where $D(k, \mu) = \det \mathbf{Q}(k, i\mu)$. For fixed $k$, these two equations define a $\mu$-parameterized hypersurface in the model's parameter space.

Instabilities with $k = 0$ are homogeneous. For $\mu = 0$, the firing rates of all units diverge. For $\mu \neq 0$, the instability leads to an oscillatory state. Instabilities with $k > 0$ correspond to the emergence of an inhomogeneous state which can be stationary ($\mu = 0$) or time dependent ($\mu \neq 0$).

Figure 3C plots the phase diagram of the model as a function of $J_{\mathrm{str,ctx}}$ and $J_{\mathrm{stn,ctx}}$ with all other parameters fixed and given Table 1. It is easy to verify that for $\mu = 0$, Eq. (13) is identically satisfied, whereas for given $k$, Eq. (13), determines $J_{\mathrm{stn,ctx}}$ as a function of $J_{\mathrm{str,ctx}}$. The pink line corresponds to this function for $k = 1$. The green line corresponds to the emergence of homogeneous oscillations; to obtain it we solved Eq. (12), Eq. (13) for $k = 0$ while varying $\mu$.

Although these calculations are in principle straightforward, the resulting expressions involve a very large number of terms. We used Mathematica [2] to handle them.

### Supplementary Figures

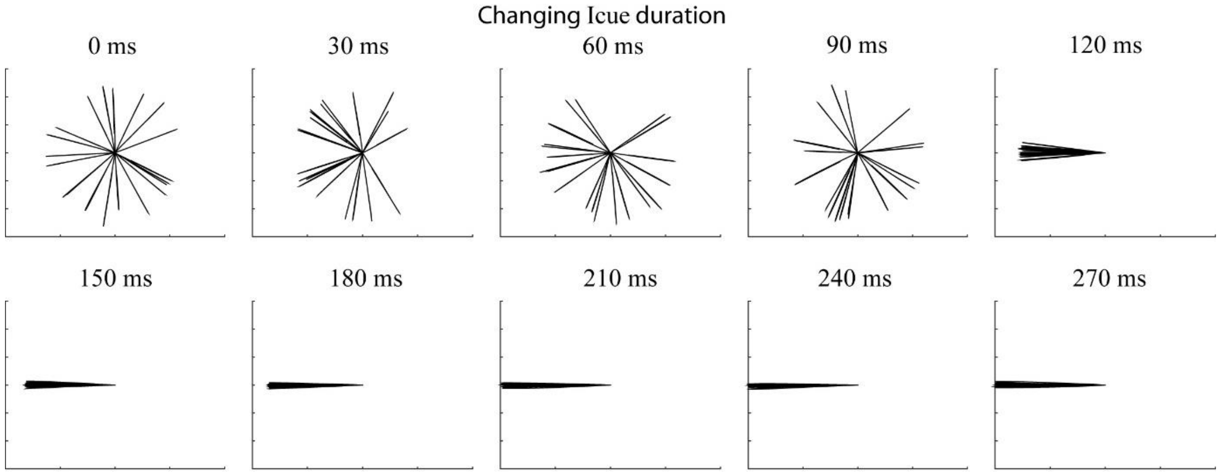


**Supplementary Figure 1:** Example trajectories (20 different renditions of a movement executed in response to a 0° cue) generated in the model for a varying duration of the input $\mathrm{I}_{\mathrm{cue}}(t)$, which is transient and shorter than the movement duration. Trajectories are computed using a population vector computed as the sum of the PD of motor cortical neurons weighted by their firing rate at each time step, as in Figure 1H of the main article. Note that the direction of the movement is well determined only if $\mathrm{I}_{\mathrm{cue}}(t)$ is sufficiently long, *i.e.*, longer than 100ms.

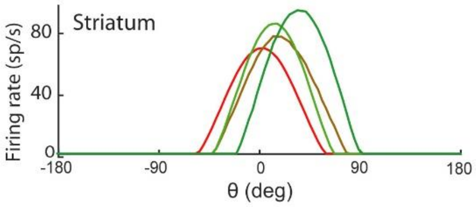


**Supplementary Figure 2:** Profile of the activity of striatal neurons across trials along the adaptation protocol (all parameters as in Figure 2 of the main paper). Example trials are coloured according to their position in the protocol (red: early, green: late). Striatal neurons receive an input from PPC (shown in Fig 2C of the main article) and an input from motor cortex. The latter shifts the peak of activity toward the movement direction. Thus, while their firing-rate profile is qualitatively similar to the input profile in Fig. 2C, it is not identical.

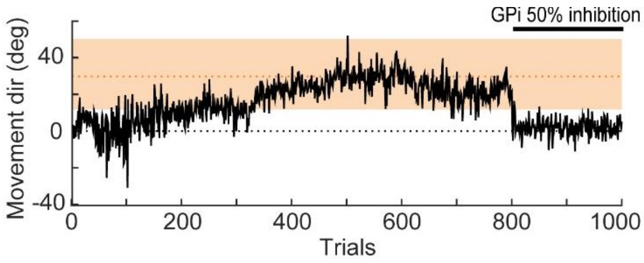


**Supplementary Figure 3:** Movement direction along the adaptation protocol in an example run with partial GPi inhibition. The orange shaded area indicates rewarded movement directions. Orange points along the x axis indicate rewarded trials. In trials 800-1000 the GPi is partially inhibited (50% inhibition).
